# Ccr4 and Pop2 control poly(A) tail length in *Saccharomyces cerevisiae*

**DOI:** 10.1101/140202

**Authors:** Vidya Balagopal, Mohan Bolisetty, Najwa Al Husaini, Jeff Coller, Brenton R. Graveley

## Abstract

Messenger RNA degradation is an important aspect of post-transcriptional gene regulation and shortening the poly(A) tail is suggested to be the rate-limiting step in mRNA degradation. In *Saccharomyces cerevisiae*, the Ccr4-Not complex is the major deadenylase and contains two subunits with exoribonuclease domains, Ccr4 and Pop2. Although the role of Ccr4 and Pop2 in deadenylation has previously been studied using individual reporter mRNAs, their activity has not been studied transcriptome-wide. Here, we describe END-seq, a method to accurately measure poly(A) tail lengths of individual mRNAs transcriptome-wide, and have used this assay to examine the impact of deleting or mutating *CCR4* and *POP2* on steady state poly(A) tail length. We found that Ccr4 and Pop2 have differential effects on the poly(A) tail lengths of individual mRNAs. Additionally, though Pop2 has previously been reported to have exonuclease activity, mutations that render it catalytically inactive have no effect on steady-state poly(A) tail lengths. Furthermore, mutations that disrupt the interaction between Ccr4 and Pop2 result in longer poly(A) tails. We also observe an inverse correlation between codon optimality and poly(A) tail length – transcripts containing predominantly optimal codons display fewer changes in poly(A) tail length upon deletion of Ccr4 or Pop2 than those containing less optimal codons. Together, these results indicate that Pop2 modulates poly(A) tail length, at least partially, via its association with Ccr4 and that Pop2 improves the function of Ccr4 in regulating poly(A) tail length. These data provide important insights into poly(A) tail length dynamics in yeast and demonstrate that END-seq is an efficient and accurate method to study poly(A) tail length.

## Introduction

Nearly all eukaryotic messenger RNAs (mRNAs) possess a 7-methylguanosine cap at the 5’ end and a poly(A) tail at the 3’ end. The 3’ ends of mRNAs are generated by co-transcriptional cleavage of the pre-mRNA and subsequent addition of non-templated adenosines by poly(A) polymerase. Poly(A) tails have been shown to play important roles in translation and RNA stability though little work has been performed to characterize poly(A) tail lengths on a transcriptome-wide scale until recently (Wiederhold and Passmore 2010; Zheng and Tian 2014). This has largely been because high-throughput sequencers have difficulties in accurately reading long homopolymers such as a poly(A) tail.

Although poly(A) tails are added to nearly all mRNAs in the nucleus during the process of 3’ end formation, poly(A) tails can also be shortened by the process of deadenylation (Parker 2012). Deadenylation is a crucial step of mRNA decay as it can remove an mRNA from the translatable pool (Garneau et al. 2007; Wiederhold and Passmore 2010). Deadenylation also initiates mRNA degradation and once the poly(A) tail is shortened to ~10-12 nucleotides (in yeast), the mRNA is decapped by Dcp1/2 and then rapidly destroyed by the 5’-to-3’ exonuclease Xrn1 (Muhlrad and Parker 1992; Decker and Parker 1993; Chen and Shyu 2011). In some cases, however, deadenylation is followed by the exosome-mediated degradation of the mRNA in the 3’-to-5’ direction (Meyer et al. 2004). Deadenylation is required for both decapping and 3’-to-5’ decay and is one of the slowest steps in the decay process making the regulation of deadenylation to be the most effective way of controlling mRNA decay (Cao and Parker 2003, 2001). Still, little is known about how poly(A) tail shortening is initiated and regulated.

In yeast and mammals, the Ccr4-Not complex has been shown to carry out the majority of cytoplasmic deadenylation (Daugeron et al. 2001; Tucker et al. 2001; Yamashita et al. 2005). The yeast Ccr4-Not complex contains Ccr4, Pop2, Not1, Not2, Not3, Not4, Not5, Caf40, and Caf130 (Denis and Chen 2003). Ccr4, a member of the endonuclease-exonuclease-phosphatase (EEP) protein family (Wang et al. 2010), and Pop2, which contains an RNase D domain from the DEDD superfamily, have been speculated to be the 3’-5’ exoribonucleases in this complex. Although Ccr4 has clearly been shown to be catalytically active in yeast, the role of Pop2 is less clear. Although it has been reported that recombinant Pop2 expressed in and purified from bacteria has exonuclease activity *in vitro* (Daugeron et al. 2001), other groups have failed to observe Pop2 exonuclease activity *in vitro* (Tucker et al. 2001) and have observed no functional impact of mutations in conserved catalytic residues of Pop2 in yeast (Chen et al. 2002). In addition, Pop2 has yeast-specific amino acid insertions in the RNase D domain that have been proposed to render the protein catalytically inactive (Tucker et al. 2001; Chen et al. 2002; Collart and Panasenko 2012). Thus, the role of Pop2 exonuclease activity in yeast remains unclear. However, the structure of the Ccr4-Not complex revealed that the interactions of Ccr4 with Pop2 and Pop2 with Not1 are important for deadenylation and decay of an *MFA2* reporter mRNA (Basquin et al. 2012).

A drawback of deadenylation studies to date is that their interpretation is often based on only a few (sometimes one) mRNAs used as reporters to assay for deadenylation. Though results obtained with single mRNAs may yield general insights, they also may not be representative of the behavior of the entire transcriptome. Most previous studies have often combined with RNase H digestion with high resolution northern blots to measure the poly(A) tail lengths of individual reporter mRNAs (Sippel et al. 1974; Decker and Parker 1993; Sheets et al. 1994). Northern blots are limited by the sensitivity and resolution of the gel and given the amount of mRNA, time and labor required for these assays, makes them unsuitable for transcriptome-wide studies.

Here, we describe a transcriptome-wide method to accurately and affordably measure poly(A) tail length. Furthermore, we deleted and made point mutations in the genes encoding Ccr4 and Pop2 and measured steady-state poly(A) tail lengths transcriptome-wide in S. *cerevisiae*. From these results, we show that deletion of each gene has substrate-specific effects on poly(A) tail length. Second, deletion of *POP2* impacts fewer mRNAs than deletion of *CCR4* and almost all Pop2 affected mRNAs overlap with those affected by Ccr4. Third, we found that while mutation of the catalytic residues of Pop2 had no detectable effect on poly(A) tail length, mutations that disrupt the Ccr4-Pop2 interaction do impact poly(A) tail lengths. These results indicate that Pop2 appears to function, at least in part, by recruiting Ccr4 and that Ccr4 might require Pop2 for efficient exonuclease activity on some mRNAs.

## Results

### Genome-wide poly(A) tail length measurements

To study the biogenesis, dynamics and function of poly(A) tail length, we designed and implemented an assay we call END-seq that accurately measures global poly(A) tail lengths (Fig. 1A). Briefly, a single round of oligo-dT selection is performed to enrich for mRNAs, an RNA linker is ligated to the 3’ end, and the RNAs are partially fragmented by limiting alkaline hydrolysis. Synthetic *in vitro* transcribed RNAs with poly(A) tails of 30 and 85 nt are added to each library after oligo-dT selection as an internal control for the accuracy of tail-length measurements. The 3’ end fragments are then selectively reverse transcribed into cDNA using a primer complementary to the linker that also contains sequences for Illumina sequencing. Second-strand synthesis is then performed using a random decamer that again contains sequences for Illumina sequencing. The double-stranded library is then amplified by PCR using primers that add Illumina flowcell binding sequences. Libraries with insert sizes ranging from 200 - 400 bp were gel purified and sequenced using paired-end sequencing on an Illumina MiSeq for 75 bp read one and 225 bp read two as well as dual indexes to sequence the barcodes that identify different libraries. The sequence of read one was aligned to the genome and used to identify the gene or transcript of origin while the sequence of read two was used to determine the length of the poly(A) tail (Fig. 1B, Materials and methods).

**Figure 1.**
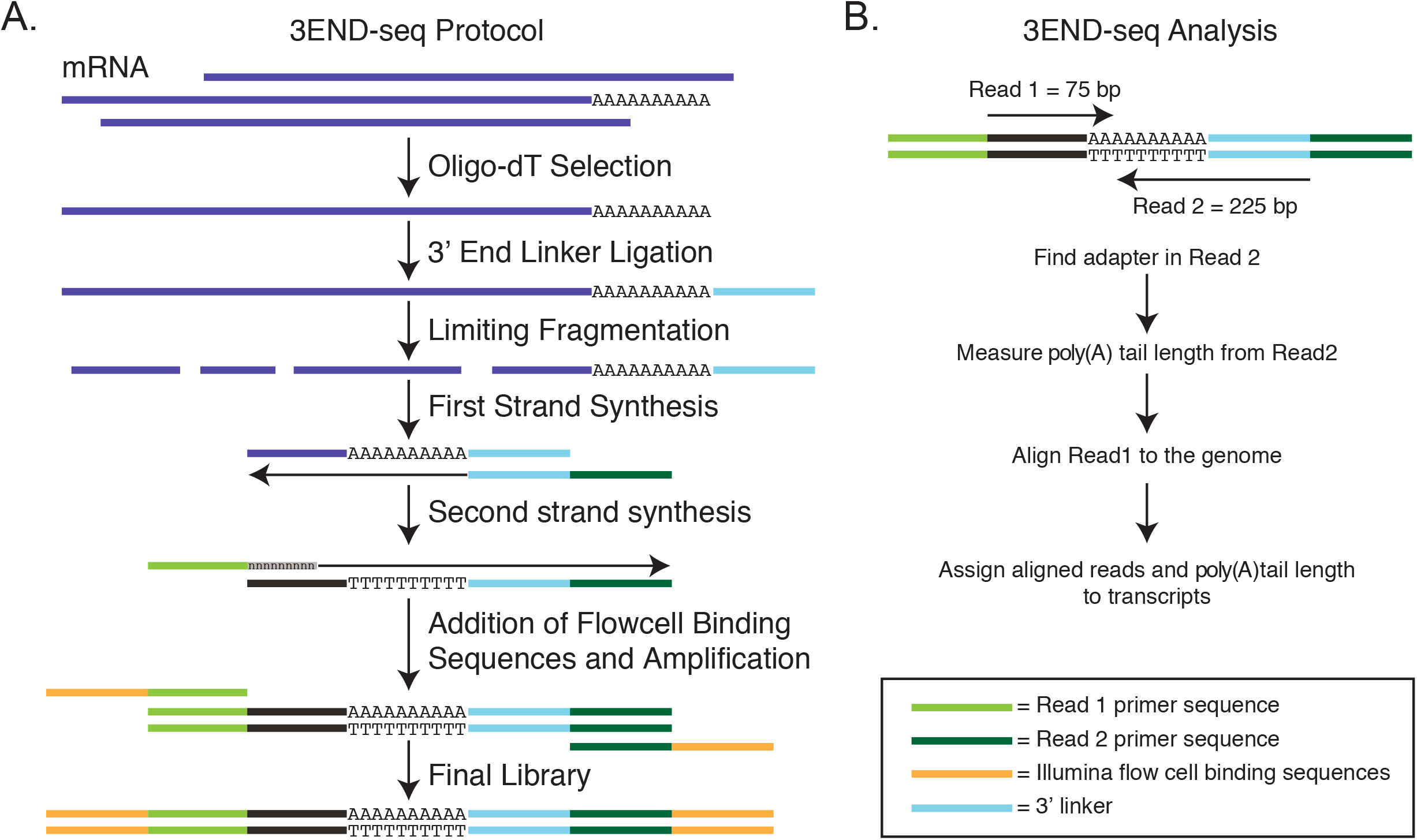
**A.** Illustration of the END-Seq protocol **B.** Schematic representation of the sequencing and analysis pipeline.

We first used END-seq to measure poly(A) tail lengths of mRNAs in wild-type (WT) S. *cerevisiae* (Table 1) grown at log phase. We obtained approximately 12 million raw read pairs for each biological replicate. These were then filtered for mate-pairs containing at least 5 Ts immediately following a perfect match to the adapter sequence present in read two. For mate-pairs that passed this filter, nucleotides 5 to 28 of read one were aligned to the S. *cerevisiae* genome and observed 4867 genes with at least one read. To have sufficient depth for statistical analyses, we only included genes with at least 30 reads for all subsequent analyses, which corresponded to 2685 genes (Table 2).

**Table 1.**
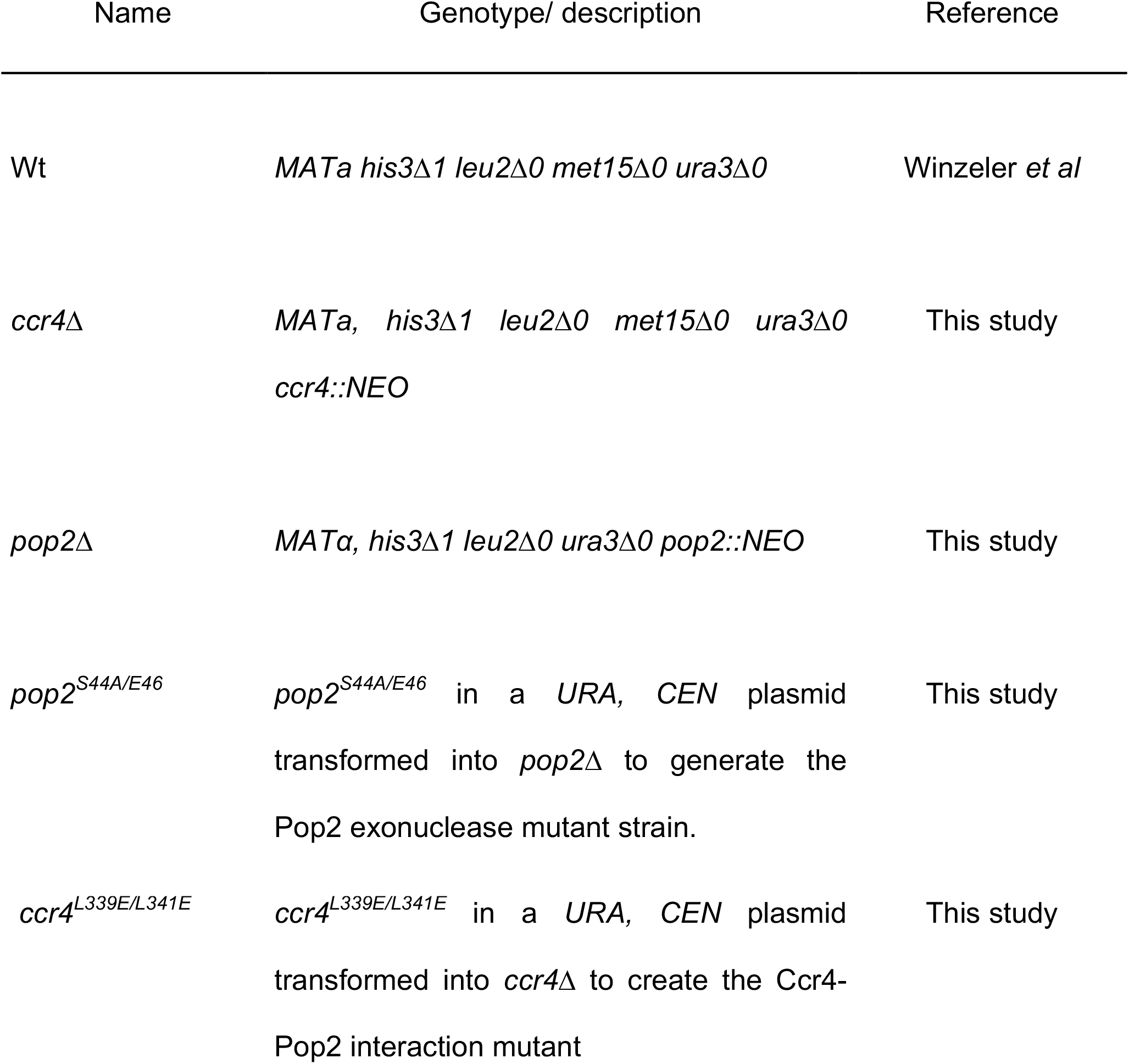
Yeast strains and plasmids used in the study.

**Table 2.** Sequencing statistics for END-seq libraries used in this study.

| **Strains** | **Total Reads** | **Reads with adapter + 5Ts** | **Reads uniquely assigned to mRNAs** | **Number of mRNAs with >30 reads** |
| --- | --- | --- | --- | --- |
| WT | 12,034,000 | 2,900,000 | 1,687,000 | 2695 |
| *ccr4*Δ | 11,610,000 | 1,870,000 | 1,170,566 | 1904 |
| *pop2*Δ | 9,052,371 | 1,904,079 | 954,331 | 2438 |
| *pop2*<sup>S44A/E46</sup> | 6,035,395 | 1,675,379 | 866,351 | 2177 |
| *ccr4*<sup>L339E/L341E</sup> | 5,549,525 | 967,759 | 492,441 | 2026 |

**Table 3.**
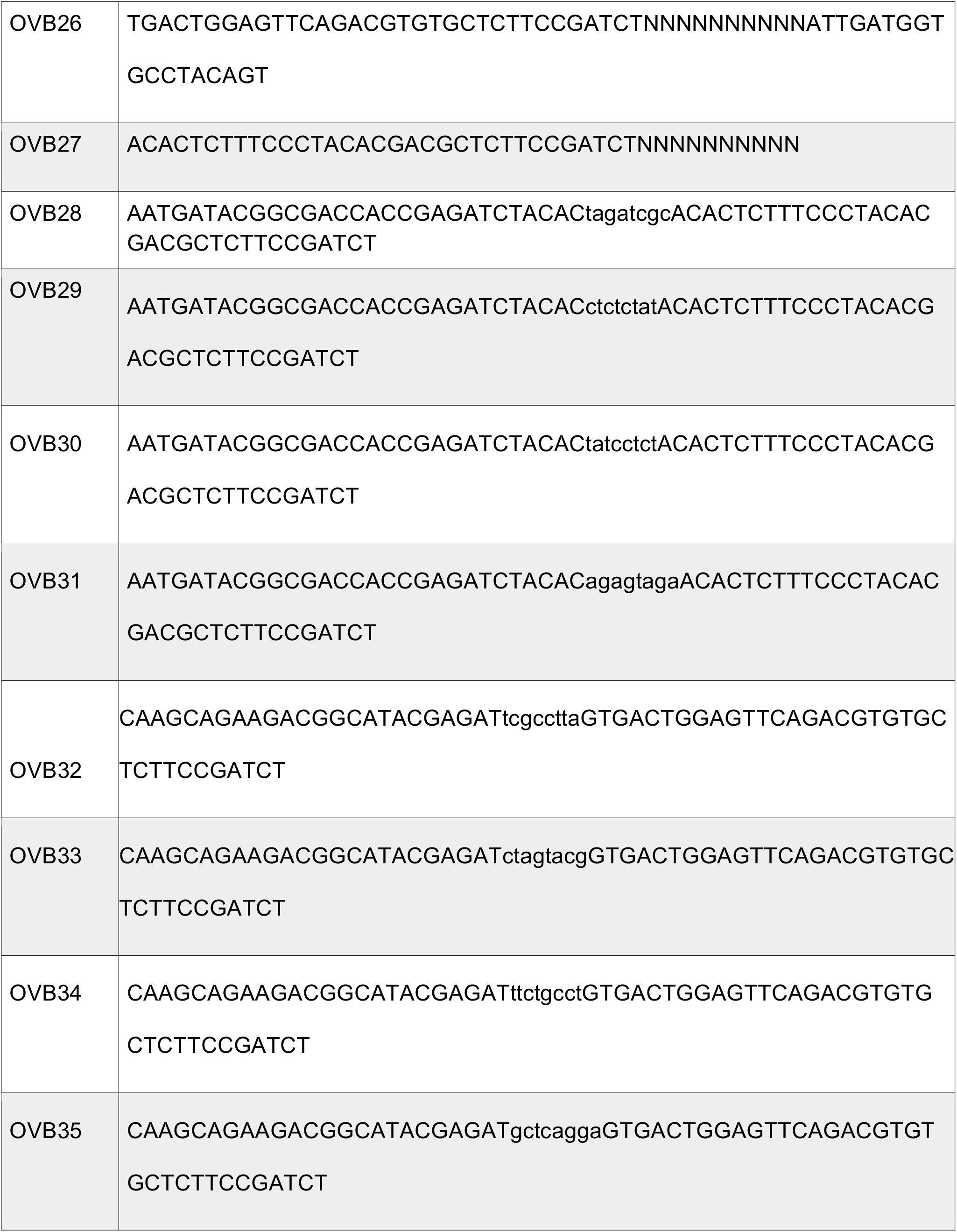
Oligonucleotides used in this study.

We performed several analyses to determine the precision and accuracy of END-seq. First, we observed a good correlation (Spearman’s r_s_ = 0.62) between RNA abundance measured by END-seq and standard poly(A)+ RNA-Seq indicating that END-seq is quantitative (Supplementary Fig. 1). Second, END-seq data is highly reproducible between biological replicates in terms of both RNA abundance and poly(A) tail length measurements, indicating high precision. Specifically, RNA abundance measurements (reads per million (RPM)) between replicates are highly correlated (Spearman’s r_s_= 0.96) (Fig. 2A). Additionally, we found the tail length distribution of all mRNAs between replicates to be nearly identical (Fig. 2B) and that the median tail lengths of individual genes is highly correlated between replicates (Spearman’s r_s_= 0.93) (Supplementary Fig. 2).

**Figure 2.**
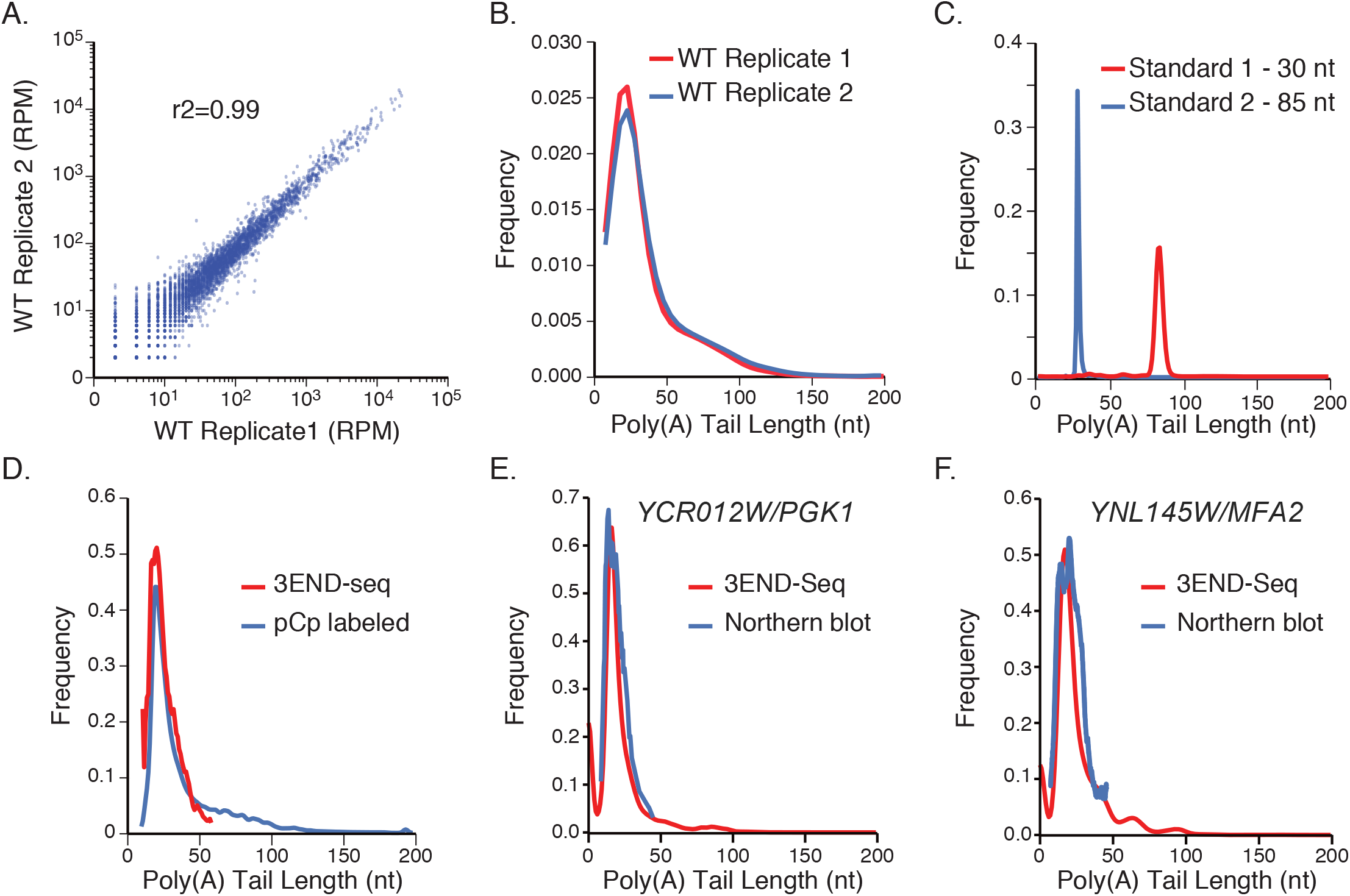
**A.** Comparison of RPM for each gene between biological replicates **B.** Transcriptome-wide poly(A) tail length measured for biological replicates is represented as a normalized histogram. **C.** Poly(A) tail length determination of two standard RNAs with poly(A) tails of known length. Two *in vitro* transcribed RNAs with known poly(A) tail lengths of 30 and 85 nt, respectively, were used as spike-ins in all END-Seq libraries. Shown are the normalized histogram plots for the poly(A) length determined for the standard RNAs using the assay and the pipeline described in Figure 1. **D.** Transcriptome-wide poly(A) tail lengths measured by END-seq is compared to global poly(A) tail measurement by pCp end labeling. **E.** Comparison of poly(A) tail length of two mRNAs *YNL145W/MFA2* and *YCR012W/PGK1* measured by northerns and END-seq.

To test the accuracy of END-seq at measuring poly(A) tail length, we examined the results obtained for the two spike-in RNAs included in each library. We found that 96% and 86% of the spike-in tail lengths were determined to be exactly 30 or 85 nt (Fig. 2C). We also compared global tail lengths determined by END-seq with bulk poly(A) tail lengths determined by pCp end labeling followed by simultaneous RNase T1 and RNase A digestion. This digestion cleaves every nucleotide but the poly(A) tract of the mRNAs and remaining poly(A) tail is resolved using denaturing polyacrylamide gel electrophoresis. We observed strong transcriptome-wide agreement between tail length measurements between these two methods (Fig. 2D). Furthermore, tail lengths measured by Northern blot analysis of the *YCR012W/PGK1* and *YNL145W/MFA2* mRNAs also agree well with those obtained by END-seq (Fig. 2E, F, Supplementary Fig. 4). These results demonstrate that END-seq provides a precise measurement of poly(A) tail length.

### Poly(A) tail length characteristics of wild-type yeast mRNA at log-phase

Our END-seq results provided tail length information for 2685 mRNAs from WT yeast at log-phase (Fig. 3A). We calculated the median tail lengths for each mRNA and found that they range from 8 nt to 105 nt with a median of 23 nt. However, mRNAs can display a wide variety of tail length distributions. For example, the *YMR044W* and *YJL134W* mRNAs both have tight mono-modal distributions, though their median tail lengths are 18 and 52 nt respectively (Fig. 3B). In contrast, *YPL122C* and *YOR293W* mRNAs have bi-modal tail length distributions (Fig. 3B). Though the underlying basis of the various modalities is unknown, they may be due to differences in the translational status, subcellular localization, or stage of the mRNA life cycle of individual mRNAs.

**Figure 3.**
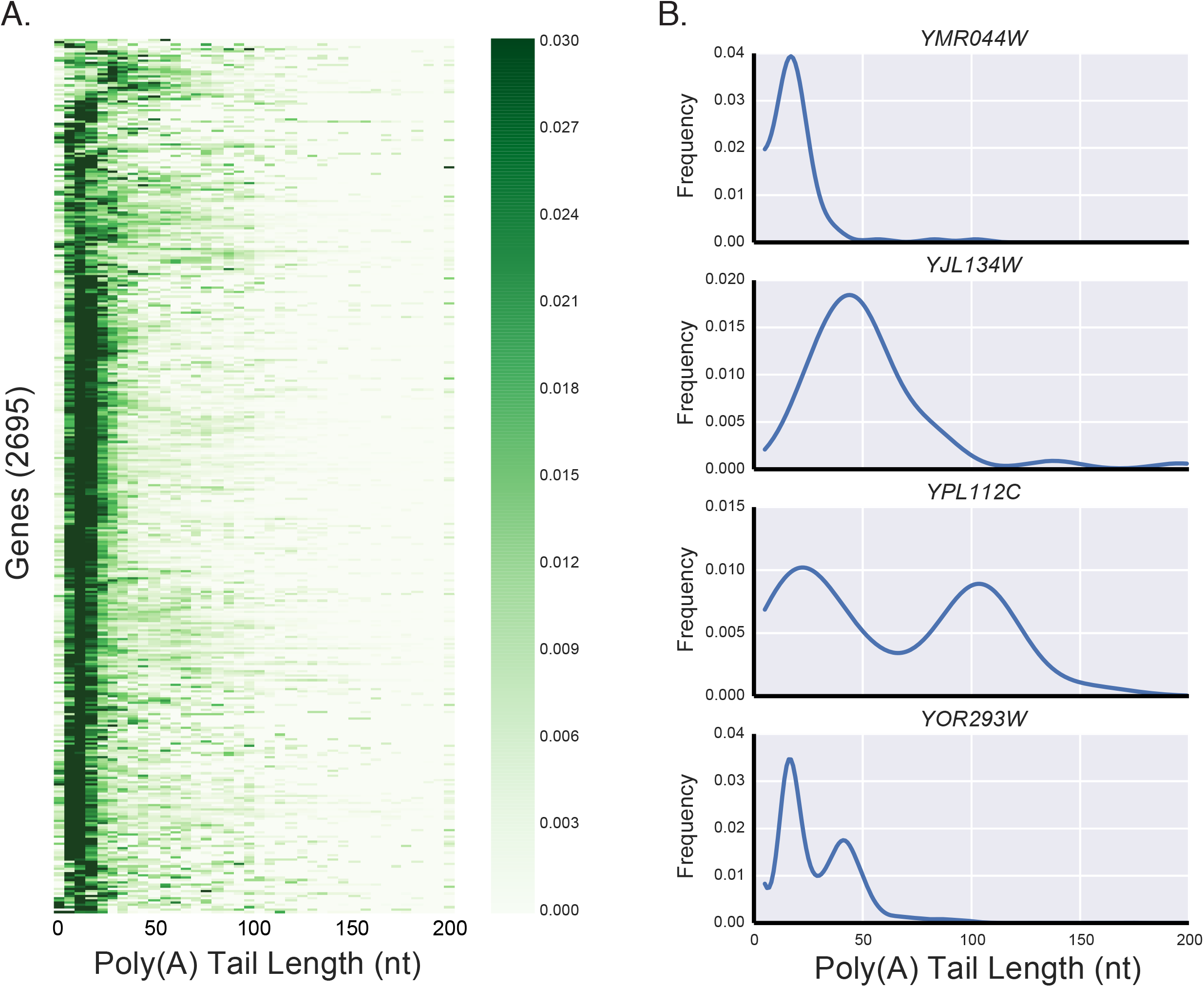
Poly(A) tail lengths of individual genes from WT yeast. **A.** Heatmap representation of the normalized poly(A) tail length distributions for individual genes. Color intensity in the heatmap indicates the fraction of reads in a bin with respect to the total reads. **B.** Normalized poly(A) tail length distributions in WT yeast for four representative mRNAs with distinct poly(A) tail length profiles.

### Ccr4 is the major deadenylase in the Ccr4-Not Complex

To identify the mRNAs whose poly(A) tail length is modulated by Ccr4 and Pop2, we used END-seq to measure poly(A) tail length transcriptome-wide at steady-state in yeast strains bearing deletions in each gene (*ccr4*Δ and *pop2*Δ). We found that the deletion of either *CCR4* or *POP2* resulted in longer poly(A) tails than in WT, both globally (Fig. 4A) and for individual mRNAs (Fig. 4B, C). It should be noted though that the poly(A) tails were longer in *ccr4*Δ cells than *pop2*Δ cells.

**Figure 4.**
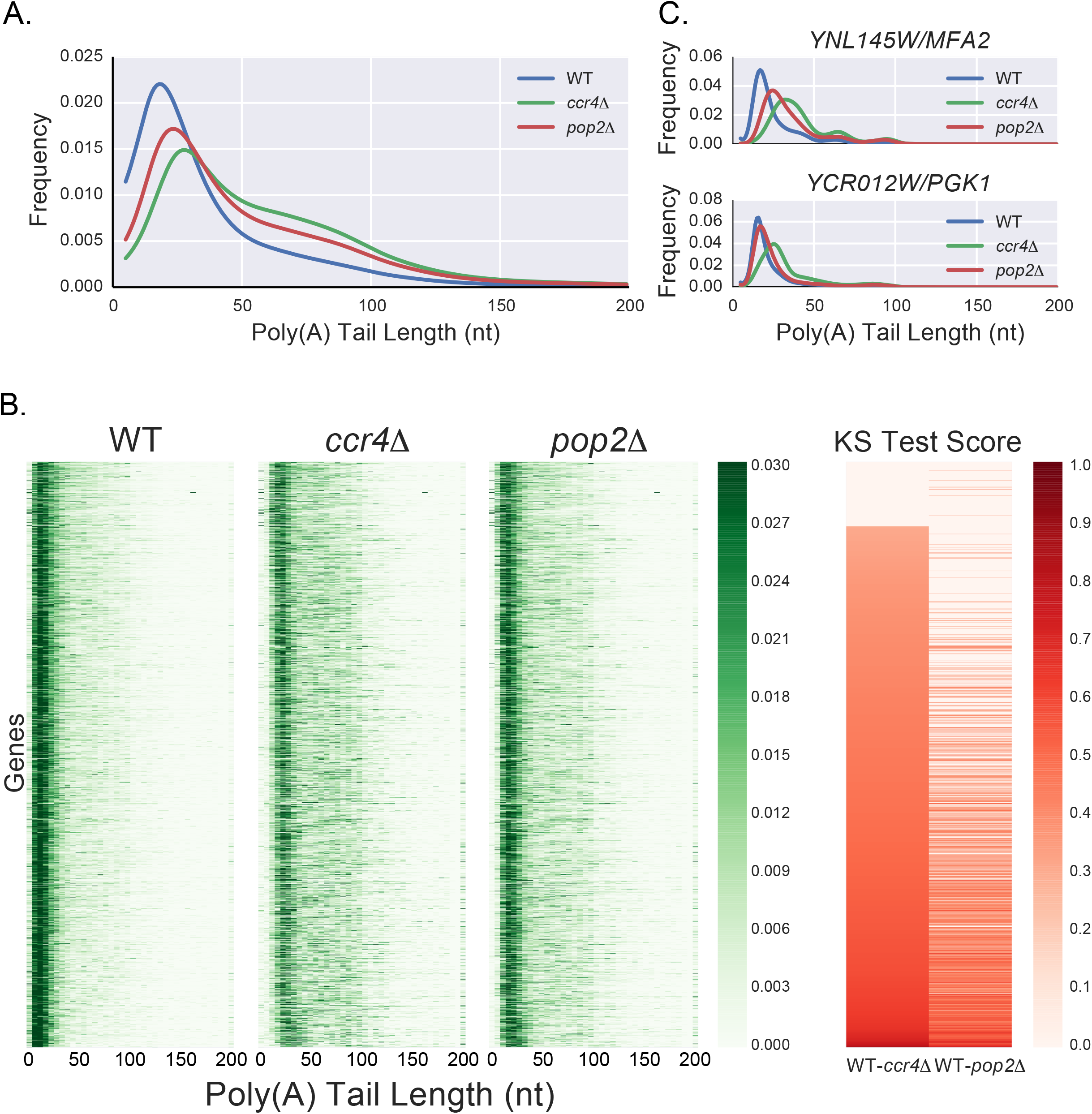
**A.** Transcriptome-wide poly(A) tail length distributions for WT, *ccr4Δ* and *pop2Δ* strains. **B.** Heatmap representation of the poly(A) tail length distributions of individual genes in WT, *ccr4Δ, pop2Δ* strains. The effect of deleting *CCR4* and *POP2* on individual mRNAs was quantitated using Kolmogorov-smirnov test scores and visualized as a heatmap. The mRNAs were sorted based on the effect of Ccr4 on poly(A) tail length of mRNA (KS test scores, D, of mRNAs *ccr4Δ* vs WT) to generate the ordering of transcripts. Each row corresponds to the same mRNA in each heatmap. All genes with a KS test score of *D*<0.3 or p-value>0.01 are depicted as white. *C*. Tail length distribution of *YNL145W/MFA2* and *YCR012W/PGK1* mRNAs in WT, *ccr4Δ, pop2Δ* strains as measured by END-seq.

We next quantitated how deletion of *CCR4* or *POP2* affected the tail length of individual genes. We used the pair-wise non-parametric Kolmogorov-Smirnov (KS) test to calculate the KS test score *(D)* to compare the tail lengths of each gene between the WT and each mutant. The value of the test score *(D)* ranges from 0 to 1. A *D* value of 0 means that distributions being compared are exactly the same while a value greater than 0 means that distributions are different from either other. As the distributions diverge, the *D* value increases to 1. Since the *D* value is an absolute value of the difference between two distributions, it does not inform about the directionality of change, i.e, longer or shorter tail lengths. Due to this we rely on the changes in median tail lengths. The difference between distributions are considered significant if the associated p-value is lower than 0.01. We found that >98% of mRNAs have *D*<0.3 between replicate experiments (Supplementary Fig. 3A) and that *D*>0.3 generally corresponds to a change in median tail length of at least 1.5-fold. We therefore used cutoffs of *D*>0.3 and p-value<0.01 for the remainder of our analyses (note: all p-values less than 0.0001 are reported as p<0.0001). To demonstrate the relationship between KS test score and poly(A) tail length change, we plotted the tail length distributions of three mRNAs, *YNL145W/MFA2, YCR012C/PGK1* and *YLR167W* in the WT, *ccr4Δ*, and *pop2Δ*. The D values and median tail lengths of these three mRNAs show a range of tail lengths and KS test scores. For example, *YNL145W/MFA2* has much longer tail lengths in *ccr4Δ* than in WT with KS test scores of *D*=0.58, p<0.0001 and median TL change of 19 nts from 18 in WT to 37 in *ccr4Δ* (Supplementary Fig. 3B). On the other hand, the tail length of *YCR012C/PGK1* is less affected than *YNL145W/MFA2* by *ccr4Δ* (*D*=0.47, p<0.0001) and median TL change of 21 from 17 in WT to 38 in *ccr4Δ* (Supplementary Fig. 3C) and finally, *YLR167W* tail lengths do not meet our D cut off of 0.3 and one sees minor changes in poly(A) tail length in *ccr4Δ* (*D*=0.28, p<0.0001). D value is more descriptive of the distribution and hence is a more accurate measure of the actual change than the median tail length.

From our data, we can draw several conclusions. First, deletion of *CCR4* altered the tail length of more mRNAs (1674 mRNAs with *D*>0.3) than the deletion of *POP2* (909 mRNAs with *D*>0.3). Interestingly nearly every mRNA whose tail length was significantly affected by deletion of *POP2* was also affected by deletion of *CCR4* (Fig 4). However, the converse is not true. For 765 mRNAs, poly(A) tail length was altered only by deletion of *CCR4* but not deletion of *POP2*.

Second, the poly(A) tail lengths of individual mRNAs were differentially affected by deletion of *CCR4* and *POP2* in most cases. This suggests that the two proteins affect the same mRNAs differently. Not all mRNAs that are impacted by the deletion of *CCR4* or *POP2* display tail length changes of the same extent (Fig. 4B). For instance, the median poly(A) tail lengths of *YCR012C/PGK1* mRNA are 17, 38 and 23 nt in WT, *ccr4*Δ (*D*=0.47, p<0.0001 *n*=8,907) and *pop2*Δ (*D*=0.19, p<0.0001, *n*=11,104), respectively. Thus, for *YCR012C/PGK1*, deadenylation is strongly impaired by deletion of *CCR4* while deletion of *POP2* has a much smaller effect (Fig. 5A). In general, deletion of *CCR4* had a greater effect on the tail length than the deletion of *POP2*. The range of change in tail length can be anywhere from unchanged to 29-fold. For instance, the median tail length of *YHL020C* mRNA changes from 5 nt in WT to 147 nt in *ccr4Δ* cells (*D*=0.87, p<0.0001, *n*=37) while the median tail length of *YPR080W* mRNA does not change in *ccr4Δ* cells (*D*=0.075, p<0.0001, *n*=7977).

**Figure 5.**
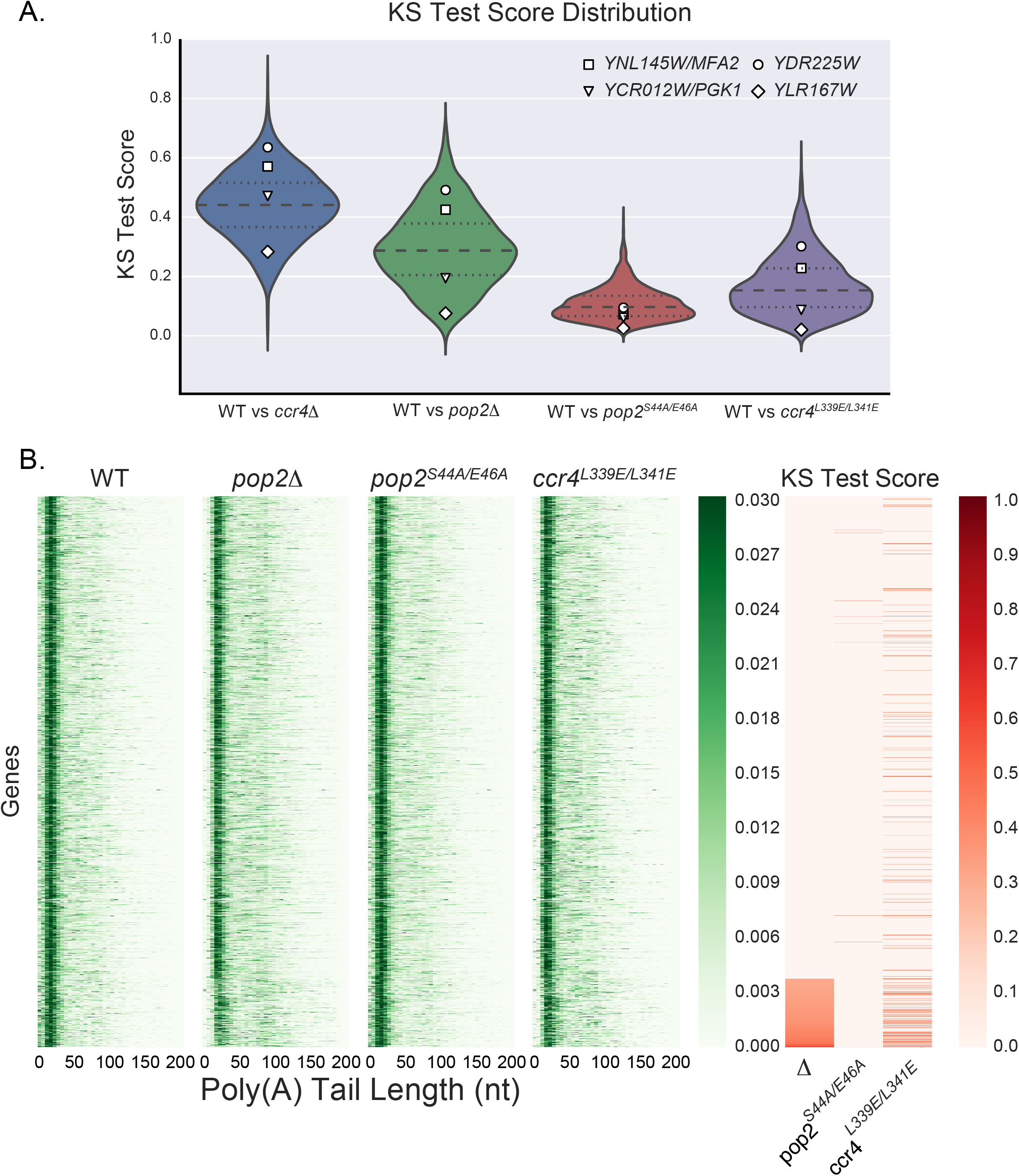
**A.** Violin plots of the distribution of KS test scores for all mRNAs studied in *ccr4Δ, pop2Δ, pop2^S44A/E46^* and ccr4^L339E/L341E^ strains. Four mRNAs *YNL145W*, *YCR012C, YDR225W* and *YLR167W* are highlighted to show the range of KS test score distributions. **B.** Heatmap of the poly(A) tail length distributions of individual genes in WT, *pop2Δ*, *pop2^S44A/E46A^*, and *ccr4^L339E/L341E^* strains. The effect of *pop2^S44A/E46A^*, and *ccr4^L339E/L341E^* point mutants on individual mRNAs was quantitated using Kolmogorov-smirnov test statistic and visualized as a heatmap. The mRNAs were sorted based on the effect of deletion of *POP2* on poly(A) tail length of mRNA (KS test score, D, of mRNAs *pop2Δ* vs WT) to generate the ordering of transcripts. Each row corresponds to the same mRNA in each heatmap. All genes with a KS test score of *D*<0.3 or p-value>0.01 are depicted as white.

**Figure 6.**
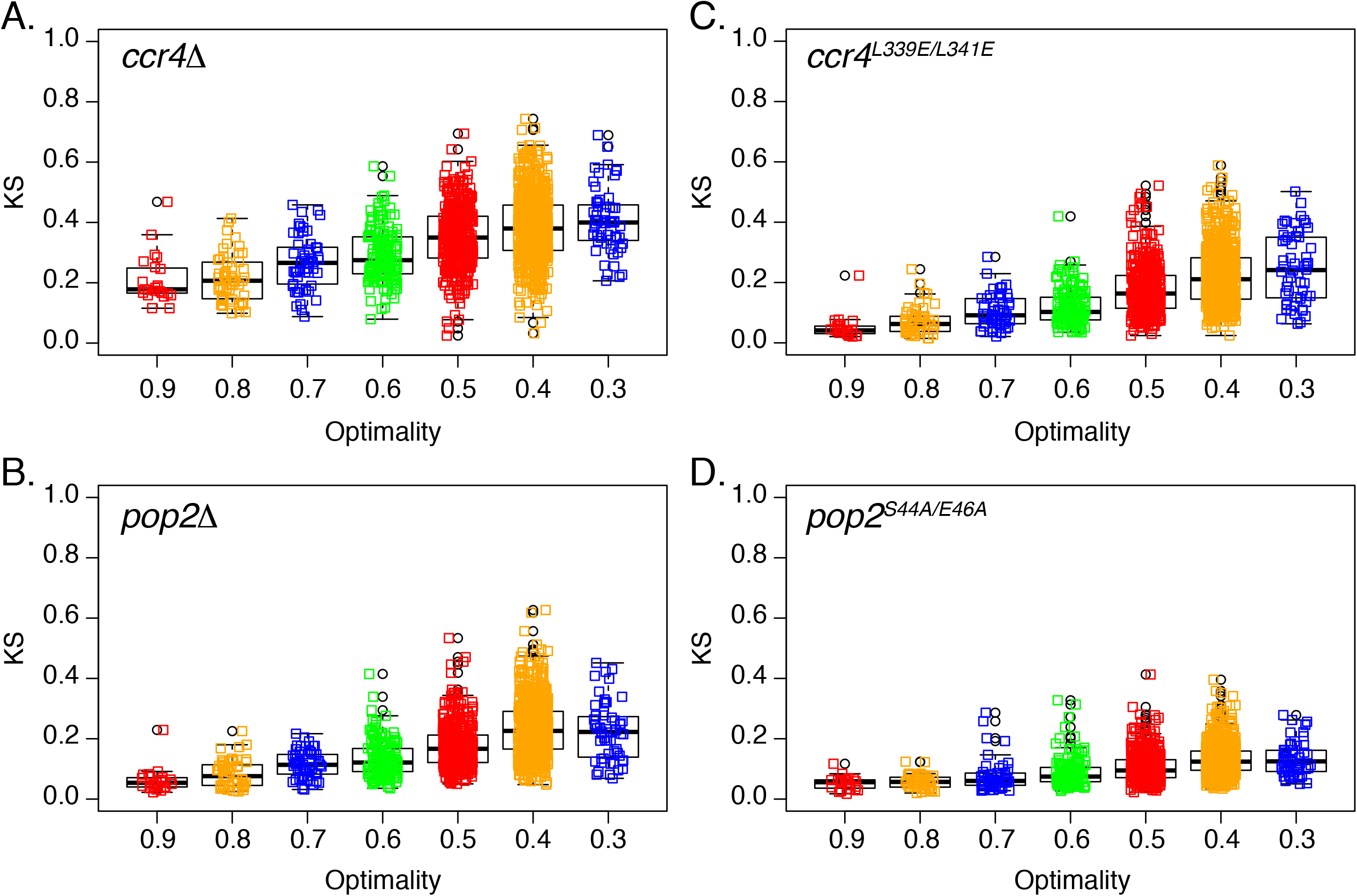
Box plot of KS test scores for all mRNAs studied in *ccr4Δ* (A), *pop2Δ*(B), *ccr4*^L339E/L341E^(C) and *pop2^S44A/E46^* (D) strains separated into optimality groups. Half of the data fall within the boxed section, with the whiskers representing the rest of the data.

There were, however 232 mRNAs, including *YNL145W/MFA2*, where both Ccr4 and Pop2 appear to play an important role in poly(A) tail length. Specifically, *YNL145W/MFA2* mRNA displayed median poly(A) tail lengths of 18, 37 and 35 nt in WT, *ccr4Δ* (D=0.57, p<0.0001, n=3,063) and *pop2Δ* (D=0.42, p<0.0001, n=4,106), respectively. Together, these results demonstrate that in most cases, Ccr4 plays a more critical role than Pop2 in determining poly(A) tail length, though for some mRNAs, Ccr4 and Pop2 have an equal effect on mRNA poly(A) tail length.

### Role of Pop2 in deadenylation

While it is clear that Ccr4 is the major deadenylase in the Ccr4-Not complex, it is less clear whether or not Pop2 has catalytic deadenylase activity *in vivo*. Nonetheless, the purified recombinant exonuclease domain of Pop2 has been shown to have exonuclease activity *in vitro* (Daugeron et al. 2001) indicating that Pop2 could be catalytically active *in vivo*. We identified 909 mRNAs whose steady-state tail lengths were significantly altered upon *POP2* deletion, all of which are affected by deletion of *CCR4* to a similar or greater extent. This result is consistent with, but does not prove that Pop2 is an exonuclease.

We propose that Pop2 either functions as an independent deadenylase or that Pop2 is not a deadenylase, but rather recruits Ccr4 which in turn deadenylates the mRNAs. To distinguish between these possibilities, we generated point mutants to either disrupt the putative exonucleolytic activity of Pop2 or disrupt the association between Ccr4 and Pop2 (Basquin et al. 2012). If Pop2 functions as an independent deadenylase, we predict that the Pop2 exonuclease mutant will have a similar affect on poly(A) tail length as a Pop2 deletion. It has been shown that simultaneous mutation of S44A and E46A abolishes Pop2 exonucleolytic activity *in vitro* (Thore et al. 2003). We therefore generated a *pop2^S44A/E46A^* yeast strain, grew it to log-phase, extracted RNA and used END-seq to measure global poly(A) tail lengths. Of the 2177 genes with at least 30 reads, only 10 showed significant changes in poly(A) tail length in *pop2^S44A/E46A^* compared to WT (*D*>0.3) (Fig. 5). However, this is unlikely to be outside of experimental or biological variation as even WT replicates show significant changes (*D*>0.3) in poly(A) tail length for 33 mRNAs. Thus, the Pop2 exonuclease mutant is indistinguishable from a WT strain in terms of poly(A) tail length of the transcriptome. This suggests that the exonucleotic activity is not essential for the function of Pop2 in modulating poly(A) tail length.

We next tested whether the interaction between Ccr4 and Pop2 is required for the role of Pop2 in modulating poly(A) tail length. The crystal structure of the Ccr4-Not complex shows that Pop2 bridges Ccr4 and the Not proteins (Basquin et al. 2012). Moreover, simultaneous mutation of both L339E and L341E in Ccr4 has been shown to abolish the interaction of Ccr4 with Pop2, but not the Pop2-Not1 interaction (Basquin et al. 2012). We therefore performed END-Seq on RNA isolated from a log-phase *ccr4^L339E/L341E^* strain we generated and identified 223 genes with significantly longer poly(A) tails in the *ccr4^L339E/L341E^* strain than in WT. This accounts for 53% (223 out of 419) of the mRNAs that display longer poly(A) tails in a *pop2Δ* strain. Furthermore, only 121 of the 223 mRNAs whose poly(A) tails were longer in the *ccr4^L339E/L341E^* strain also had longer poly(A) tails in the *pop2Δ* strain (Fig. 5). Interestingly, all of the 223 mRNAs with longer poly(A) tails in the *ccr4^L339E/L341E^* strain were also longer in the *ccr4Δ* strain (Fig 5). These results indicate that Pop2 participates in modulating poly(A) tail length *in vivo* by recruiting Ccr4 to specific mRNAs but that it is Ccr4 that catalyzes the deadenylation of these mRNAs.

### Codon optimality and Tail length

Our laboratories previously showed that codon optimality is a major determinant of mRNA decay in S. *cerevisiae* (Presnyak et al. 2015). Specifically, “optimal” codons, that are decoded efficiently during translation, are associated with increased mRNA stability, while “non-optimal” codons, which are decoded slowly, are associated with decreased mRNA stability. Subsequently, several other studies demonstrated that codon-mediated regulation of mRNA stability also occurs in *Escherichia coli, S. pombe, D. rerio, D. melanogaster, X. laevis*, and *M. musculus* suggesting that this is a highly conserved phenomenon (Boël et al. 2016; Harigaya and Parker 2016; Mishima and Tomari 2016; Bazzini et al. 2016). In zebrafish, two groups demonstrated that maternal mRNAs with lower codon optimality have shorter poly(A) tails and the shortening of poly(A) tails is mediated by the activity of the CCR4–NOT complex (Bazzini et al. 2016; Mishima and Tomari 2016).

We therefore explored whether there was a correlation between codon optimality and the observed effects of CCR4 and POP2 deletion or mutation on poly(A) tail length. In each case, we found that transcripts with the lowest codon optimality scores were most susceptible to changes in poly(A) tail length (Fig. 4). For example, upon deletion of *CCR4*, transcripts with an optimality score of 0.3 had an average KS value of 0.4, while those with an optimality score of 0.9 had an average KS value of only 0.2 (Fig. 4A). These differences are most pronounced in the *ccr4Δ* strain (Fig. 4A), which displayed the greatest change in poly(A) tail length and lowest in the *pop2^S44A/E46A^* strain nwhich had the smallest differences in poly(A) tail length (Fig. 4D). Thus, these results indicate a close relationship between codon optimality, RNA stability, and poly(A) tail length.

## Discussion

This transcriptome-wide study provides several important insights into how Ccr4 and Pop2 modulate poly(A) tail length in S. *cerevisiae*. First, we identified the mRNAs whose poly(A) tail length is altered by deletion of *CCR4* and *POP2* and show that these proteins have differential affects on poly(A) tail length of different mRNAs. Second, Pop2 modulates the poly(A) tail length of fewer mRNAs than Ccr4, and almost all Pop2 targets are also Ccr4 targets, though many Ccr4 targets are not targets of Pop2. Finally, Pop2 exonuclease activity is not required to modulate poly(A) tail length *in vivo*, but the interaction between Pop2 and Ccr4 is required for proper poly(A) tail maintenance of several mRNA. This suggests that rather than deadenylating RNAs directly, Pop2 functions by recruiting Ccr4 to Pop2-dependent mRNAs and that it is Ccr4 that deadenylates these mRNAs.

## Poly(A) tail length measurement assays

Poly(A) tail length is both an important and understudied aspect of gene regulation. The main reason this topic has not been extensively studied is simply due to the lack of assays that can monitor poly(A) tail length genome-wide, and this motivated our effort to develop such an assay. While this work was in progress two papers were published describing techniques to measure poly(A) tail length genome wide (Chang et al. 2014; Subtelny et al. 2014). PAL-seq used a modified Illumina Genome Analyzer II (GAII) sequencer to measure poly(A) tail length based on the fluorescence intensity of probe hybridization to poly(A) tails (Subtelny et al. 2014). Though powerful, PAL-seq has two limitations. First, PAL-seq requires access to and extensive modification of an Illumina GAII to perform this assay. Second, PAL-seq uses fluorescence intensity to measure poly(A) tail length and accordingly, PAL-seq has a resolution of +/− 25 nt. In contrast, END-seq has single nucleotide resolution. Nonetheless, the poly(A) tail length distributions determined by PAL-seq and END-seq for WT yeast is comparable. Specifically, the global median poly(A) tail length measured by PAL-seq is 26.3 nt compared to 23 nt when measured by END-seq

In contrast to PAL-seq, TAIL-seq (Chang et al. 2014; Lim et al. 2016) is very similar to END-seq. The major differences between TAIL-seq and END-seq is that TAIL-seq uses enzymatic fragmentation of the RNA and the ligation of RNA linkers to both the 3’ and 5’ ends of the mRNA while END-seq uses alkaline hydrolysis and the ligation of an RNA linker to the only the 3’ end followed by random decamer primed second strand synthesis. Unfortunately, as only human poly(A) tail lengths were determined with TAIL-seq, we are unable to directly compare the accuracy and performance of these two assays. Nonetheless, given the similarity of the assays we expect that they should perform similarly. We anticipate that the development of these new assays that accurately measure poly(A) tail length will be used to provide new insights into the role of poly(A) tail length in biology.

## Role of Ccr4 and Pop2 in deadenylation

This study has identified the impact of Ccr4 and Pop2 on poly(A) tail length at logphase in yeast and confirms the varying effect of these two proteins on poly(A) tail length. Ccr4 appears to be a stronger deadenylase than Pop2. This conclusion is supported by several observations. First, Deletion of *CCR4* affects the poly(A) tail lengths of substantially more genes than *POP2*. Additionally, the bulk median poly(A) tail lengths observed in WT, *ccr4Δ* and *pop2Δ* strains are 22, 35 and 28 nt, respectively. Finally, Ccr4 and Pop2 both affect the poly(A) tail lengths of individual genes differently. Taken together these data demonstrate that different mRNAs have different deadenylation rates and that these dynamics could be an important point of regulation of gene expression by controlling mRNA stability.

A much debated topic is the function of Pop2 in the Ccr4 complex. Even though Ccr4 and Pop2 are both part of the Ccr4-Not complex, deletion of *CCR4* has a greater effect on poly(A) tail length than deleting *POP2*, both in terms of the number of mRNAs affected as well as the magnitude of increase in the poly(A) tail length. Additionally, all mRNA whose poly(A) tail length is affected by deletion of *POP2* are also equally or more strongly affected by deletion of *CCR4*. This suggests that Pop2 function requires Ccr4. For example, Pop2 may have minimal deadenylase activity that is only active in the presence of Ccr4. Alternatively, Pop2 may not have deadenylase activity itself, but rather functions to promote the deadenylase activity of Ccr4. This is supported by our observation that poly(A) tail length was essentially unchanged in the *pop2^S44A/E46A^* strain, which expresses a catalytically inactive form of Pop2. On the other hand, we observed substantial changes to poly(A) tail length in the *ccr4^L339E/L341E^* strain, which expresses a form of Ccr4 that is unable to interact with Pop2. Pop2 could function to increase the activity of Ccr4 by either recruiting Ccr4 to specific mRNAs or by increasing the processivity of Ccr4.

Previous work from our labs and others have demonstrated the correlation between codon optimality and mRNA decay rates (Presnyak et al. 2015; Boël et al. 2016; Harigaya and Parker 2016; Mishima and Tomari 2016; Bazzini et al. 2016). This study provides a possible mechanistic link between these two measures. Data from this study suggests that mRNAs with low codon optimality are more susceptible to *CCR4 and POP2* mediated deadenylation and hence more prone to decay. Recent reports have shown very weak correlations between steady-state poly(A) tail lengths and mRNA decay rates (Subtelny et al. 2014). Our data suggests that it is not the steady-state length of the poly(A) tail but rather its susceptibility to deadenylation that correlates with codon optimality and mRNA decay rates.

To asses this, we compared the effect of deletion *CCR4* and *POP2* on poly(A) tail length to the decay rates of individual mRNAs obtained from published studies (Miller et al. 2011). Changes in poly(A) tail length caused by deletion of both *CCR4* and *POP2* show a positive correlation with decay (*ccr4*Δ r_s_=0.4 and *pop2*Δ r_s_=0.6). Even though deletion of *CCR4* has a greater effect on poly(A) tail length than *POP2*, tail length changes observed upon deletion of *POP2* are more correlated with mRNA decay rates. A subset of mRNAs whose tail length is equally affected by both Ccr4 and Pop2 are the most rapidly degraded, while mRNAs whose tail length is least affected by Pop2 have very long half-lives. These results lead us to speculate that Pop2 might have a function in promoting decay.

Pop2 is known to link Ccr4 and Not proteins and hence stabilize the Ccr4-Not complex. This could in turn improve function of Ccr4 leading to faster deadenylation resulting in faster decay. It has also recently been shown that Not2, Not3 and Not5 participate in decapping potentially via interactions with the decapping activator protein Pat1 (Alhusaini and Coller 2016). Pop2 could also promote decay by recruiting decay machinery to the mRNA. Affinity capture experiments and two-hybrid assays have shown that Pop2 protein associates with Dhh1, a known activator of decapping (Hata et al. 1998; Cheng et al. 2005; Coller et al. 2001).

## Acknowledgements

We thank all members of the Graveley lab, especially Mike Duff, Alex Plocik and Sandy Garett for helpful discussions, technical assistance and critical review of the manuscript. This work was supported by funds from grants R01GM080465 to J.C. and R35GM118140 to B.R.G. M.B. was funded by AHA founders affiliate postdoctoral fellowship grant 14POST18750000.

## Materials and Methods

### Yeast strains, oligonucleotides and growth conditions

The genotypes of all strains and plasmids used in this study are listed in Table 1. Strains were grown in standard yeast extract/peptone medium (YP) or minimal media with the addition of appropriate amino acids and 2% dextrose at 30°C to mid-log phase 0.3-0.4 OD. Oligonucleotides used in the study are listed in Table 2.

### RNA analysis

RNA isolation, Northern analysis, Transcriptional shut-off experiments were performed as described in (Coller and Parker 2005). Bulk mRNA tail measurements using pCp end labeling was conducted as preciously described in (Minvielle-Sebastia et al. 1991).

### END-seq protocol

From 10 μg of total RNA, mRNA was enriched using oligo(dT) Dynabeads (Invitrogen) following the manufacturers protocol. 100 pg of IDT1 linker was ligated to mRNA using 200 units of RNA ligase 2, truncated (NEB) for 2 hrs at 25°C. Ligated mRNA was precipitated using isopropanol and glycoblue at -20°C overnight. mRNA was then subjected to partial alkaline hydrolysis in 1X RNA fragmentation buffer (Ambion) at 70°C for 1 min and stopped by the addition of EDTA to a final concentration of 0.05 M. Excess linker and short fragments were removed using Agencourt RNAClean XP following manufacturer’s protocol. cDNA was synthesized using oligonucleotide OVB26 complementary to the linker at 42°C for 18 mins. After cDNA synthesis was complete, RNA was destroyed by the addition of NaOH to a final concentration of 0.1N and heating to 98°C for 20 mins. After neutralization of the NaOH with equimolar HCl, Agencourt RNAClean XP was once again employed to remove excess oligos. Second strand synthesis was performed using oligonucleotide OVB27 using Klenow at 25°C for 2 hrs. Ampure XP cleanup was performed to remove excess oligonucleotide OVB27. Library was amplified by 14 cycles of PCR using the amplification primers (OVB 28 - OVB36) at 58°C annealing temperature and 25 seconds of extension at 68°C. Libraries were run on a 6% polyacrylamide gel and fragments in the range of 300 to 500 bp purified. DNA was eluted from the gel as described in Ingolia *et al* (Ingolia 2010).

### Quantitation of Libraries and Sequencing

Libraries were quantitated using an Agilent Tape station and a Qubit. Three libraries at 2 nM final concentration were multiplexed on a single Illumina MiSeq flow cell along with 15% phiX and sequenced to generate 75 bp from read one and 225 bp from read two, along with an 8 bp index read to deconvolute the samples.

### END-seq analysis pipeline

Read one was used to identify the gene from which the poly(A) tail was derived. 25 nt sequence starting at the 5th base of read one was aligned to the SacCer2 genome using Bowtie version 0.12.7. Only unique alignments were considered for further analysis. The alignments were then assigned to an mRNA isoform using custom scripts. All scripts used for analysis and figures in this study can be obtained on github at https://github.com/mohanbolisetty/endseq.

### Tail length measurement

Our initial attempts to measure poly(A) tail length by simply identifying the RNA linker and then counting the number of T’s proved ineffective. We determined that the base calling software would introduce non-T bases in the middle of long T runs in read two. To circumvent we tried to determine the tail length by three different methods. First, simple counting with an error rate of 10, counting the number of T’s in a sliding window of 10 nucleotides. Second, when the number of T’s in a window of 10 falls below 5, it was called the end of the tail length. Third, two strings of 6 Ts with 20 bases in between. The bases in between may contain any percent of Ts. We used the standard RNA spike-ins to test which of these measurements showed highest accuracy. The string counting was found to be closest to the actual Tail length.

## Supplementary Figure Legends

**Supplementary Figure 1.**
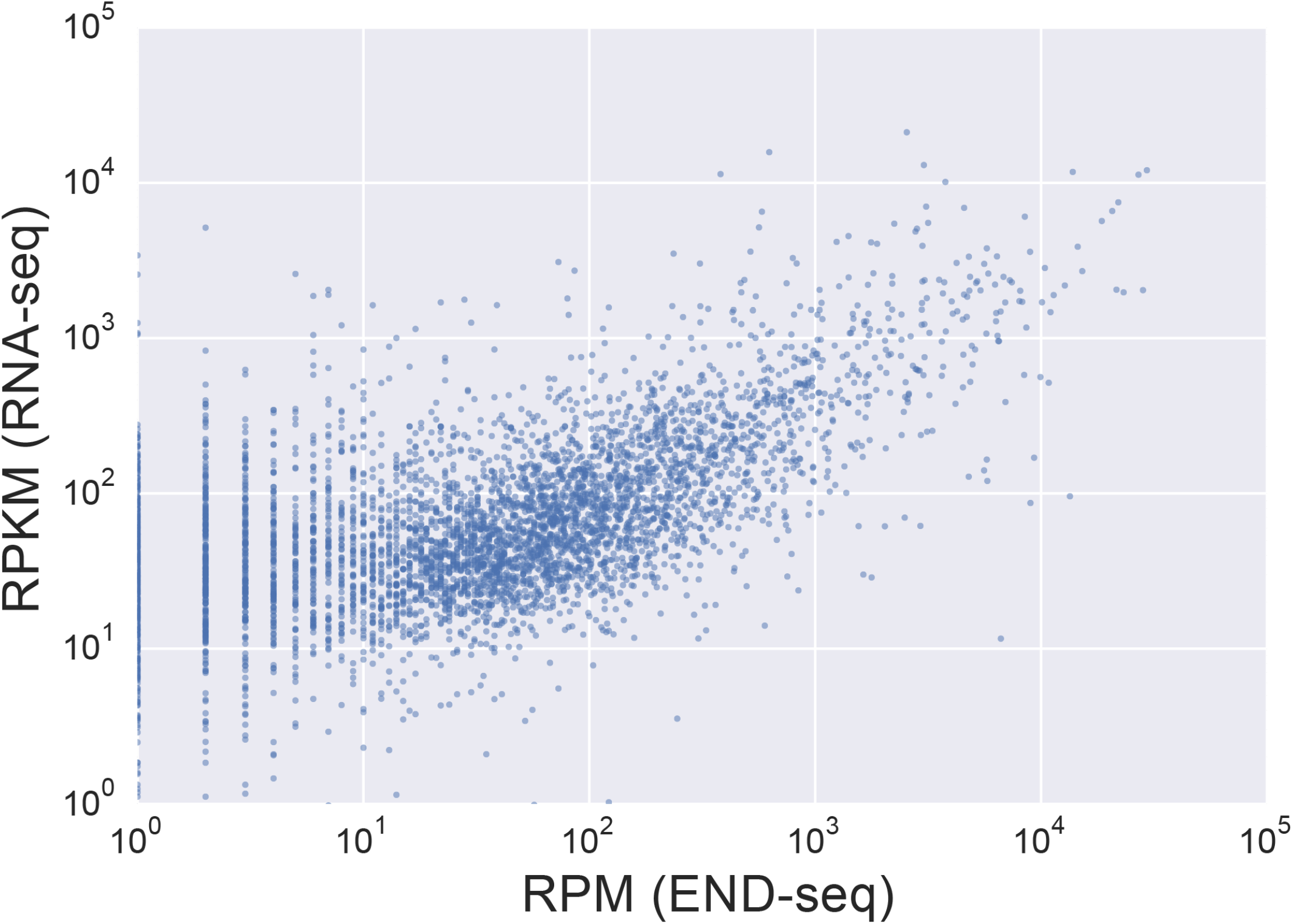
Scatter plot comparing RNA-seq (RPKM) and END-seq (RPM) from WT yeast strains.

**Supplementary Figure 2.**
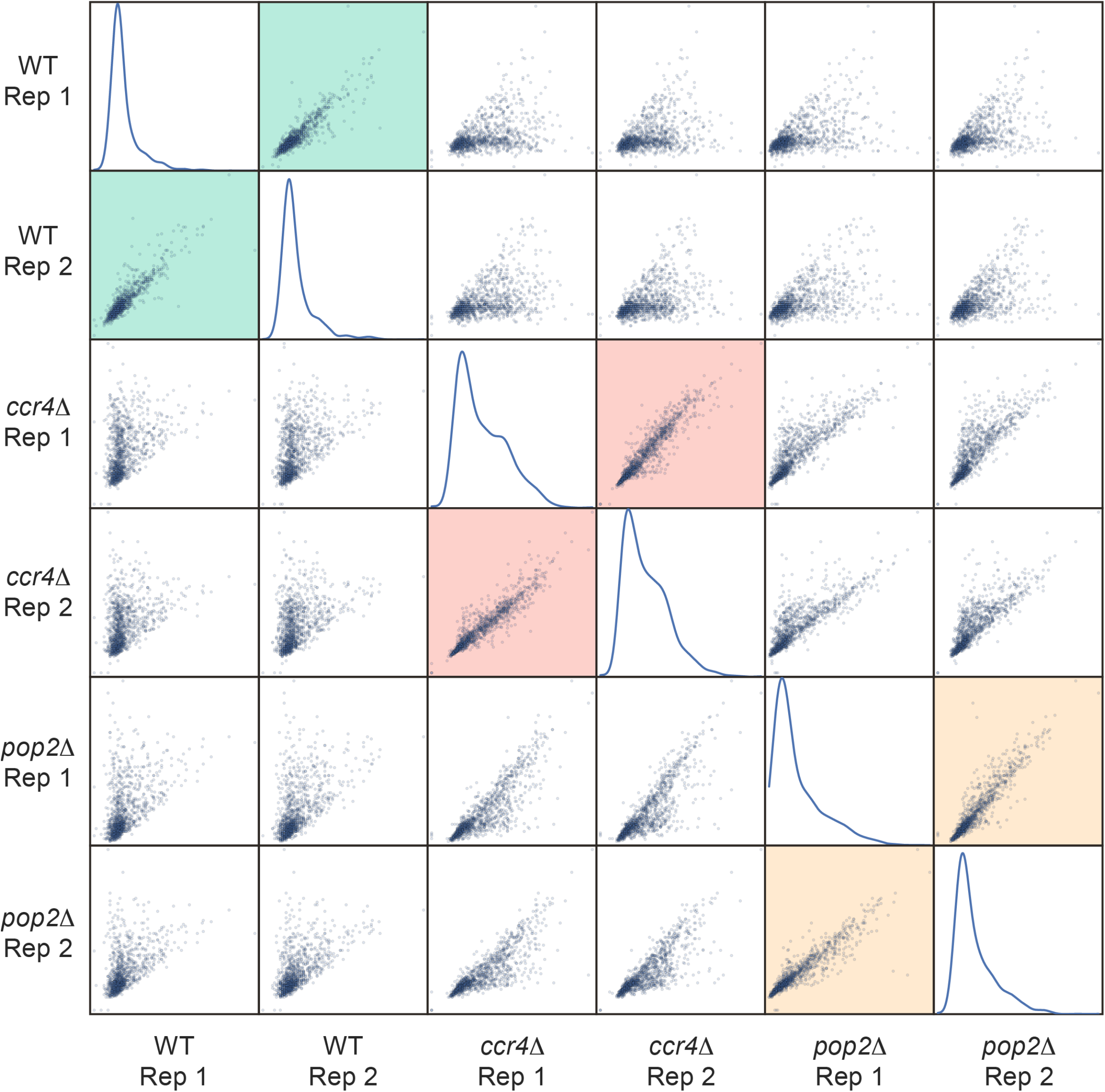
Scatter plots of the median tail length obtained the replicates of the WT, *ccr4Δ*, and *pop2Δ* strains subjected to END-seq.

**Supplementary Figure 3.**
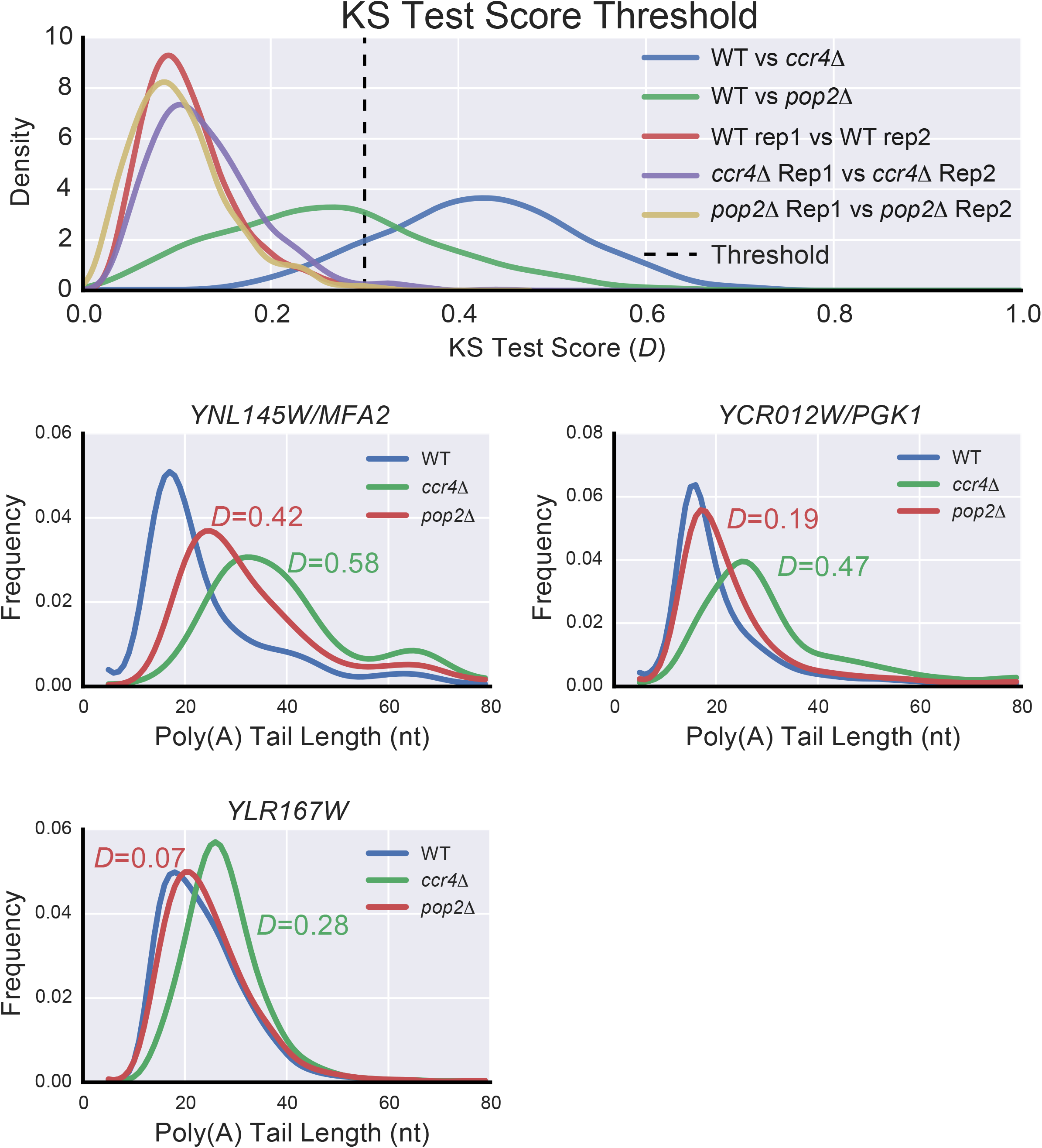
**A.** KS test score distribution for WT replicate1 vs WT replicate2, *ccr4Δ* replicate 1 vs *ccr4Δ* replicate 2, *pop2Δ* replicate1 vs *pop2Δ* replicate 2, WT vs *ccr4Δ*, and WT vs *pop2Δ*. In every the replicate comparison, >98% of the mRNAs have KS test scores *D*<0.3. **B.** Shown are the poly(A) tail length distribution of three mRNAs, *YNL145W, YCR012C* and *YLR167W*, in WT, *ccr4Δ* and *pop2Δ* strains. These mRNAs show a range of KS test scores.

**Supplementary Figure 4.**
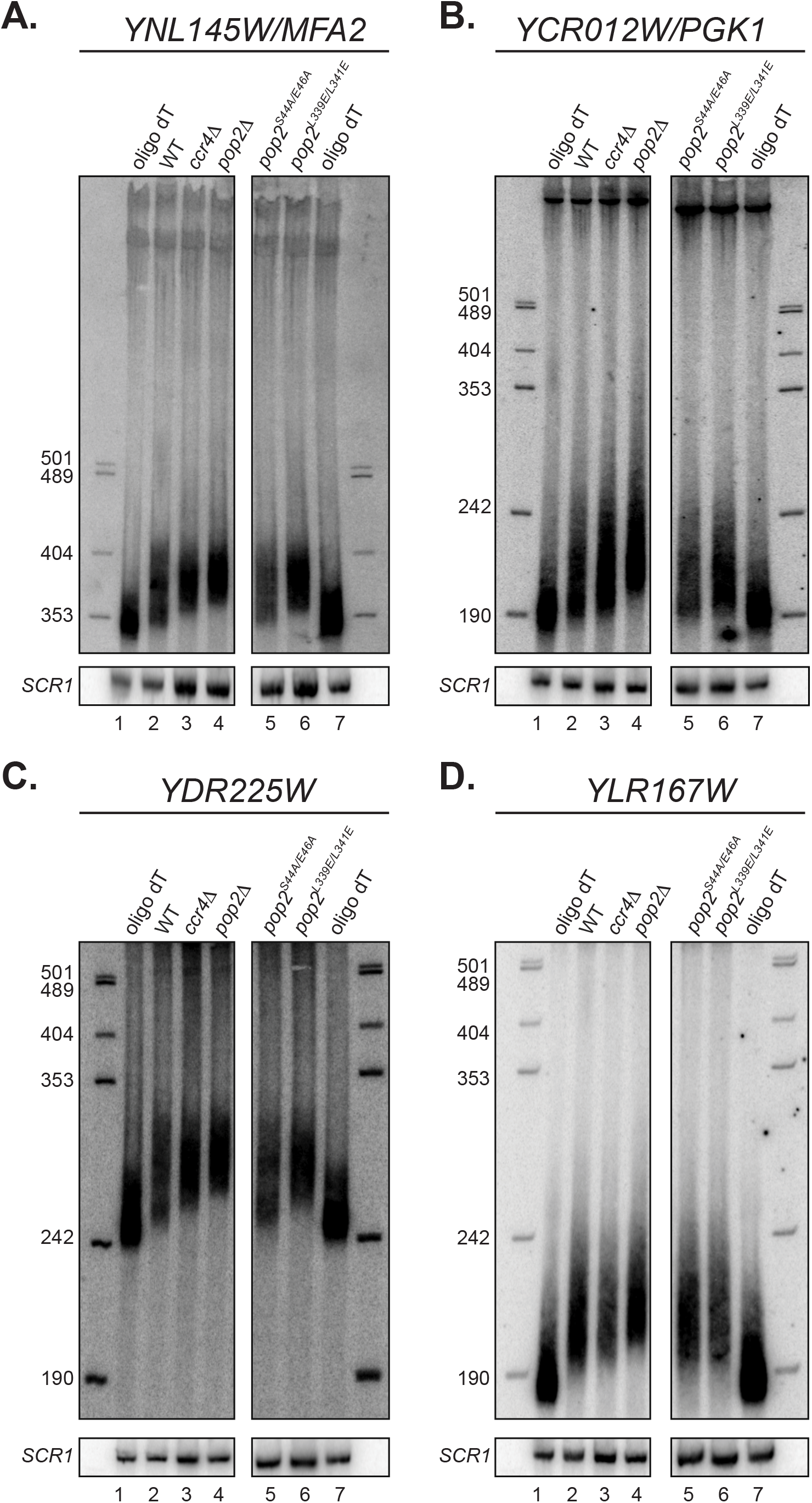
Northern analysis validation of poly(A) tail length for the four mRNAs **(A)** *YNL145W*, **(B)** *YCR012C*, **(C)** *YDR225W* and **(D)** *YLR167W* in WT, *ccr4Δ, pop2Δ, pop2^S44A/E46^* and *ccr4^L339E/L341E^*strains.

## Reference

Alhusaini N, Coller J. 2016. The deadenylase components Not2p, Not3p, and Not5p promote mRNA decapping. RNA 22: 709–21.

Basquin J, Roudko V V., Rode M, Basquin C, Séraphin B, Conti E. 2012. Architecture of the nuclease module of the yeast Ccr4-Not complex: The Not1-Caf1-Ccr4 interaction. Mol Cell 48: 207–218.

Bazzini AA, del Viso F, Moreno-Mateos MA, Johnstone TG, Vejnar CE, Qin Y, Yao J, Khokha MK, Giraldez AJ. 2016. Codon identity regulates mRNA stability and translation efficiency during the maternal-to-zygotic transition. EMBO J 35: 1721–1843.

Boёl G, Letso R, Neely H, Price WN, Wong K, Su M, Luff JD, Valecha M, Everett JK, Acton TB, et al. 2016. Codon influence on protein expression in E. coli correlates with mRNA levels. Nature 529: 358–363.

Cao D, Parker R. 2003. Computational modeling and experimental analysis of nonsense-mediated decay in yeast. Cell 113: 533–545.

Cao D, Parker R. 2001. Computational modeling of eukaryotic mRNA turnover. RNA 7: 1192–212.

Chang H, Lim J, Ha M, Kim VN. 2014. TAIL-seq: Genome-wide determination of poly(A) tail length and 3’ end modifications. Mol Cell 53: 1044–1052.

Chen CYA, Shyu A Bin. 2011. Mechanisms of deadenylation-dependent decay. Wiley Interdiscip Rev RNA 2: 167–183.

Chen J, Chiang YC, Denis CL. 2002. CCR4, a 3’-5’ poly(A) RNA and ssDNA exonuclease, is the catalytic component of the cytoplasmic deadenylase. EMBO J 21: 1414–1426.

Cheng Z, Coller J, Parker R, Song H. 2005. Crystal structure and functional analysis of DEAD-box protein Dhh1p. RNA 11: 1258–1270.

Collart MA, Panasenko OO. 2012. The Ccr4-Not complex. Gene 492: 42–53.

Coller J, Parker R. 2005. General translational repression by activators of mRNA decapping. Cell 122: 875–886.

Coller JM, Tucker M, Sheth U, Valencia-Sanchez MA, Parker R. 2001. The DEAD box helicase, Dhh1p, functions in mRNA decapping and interacts with both the decapping and deadenylase complexes. RNA 7: 1717–27.

Daugeron MC, Mauxion F, Séraphin B. 2001. The yeast POP2 gene encodes a nuclease involved in mRNA deadenylation. Nucleic Acids Res 29: 2448–2455.

Decker CJ, Parker R. 1993. A turnover pathway for both stable and unstable mRNAs in yeast: Evidence for a requirement for deadenylation. Genes Dev 7: 1632–1643.

Denis CL, Chen J. 2003. The CCR4-NOT Complex Plays Diverse Roles in mRNA Metabolism. Prog Nucleic Acid Res Mol Biol 73: 221–250.

Garneau NL, Wilusz J, Wilusz CJ. 2007. The highways and byways of mRNA decay. Nat Rev Mol Cell Biol 8: 113–126.

Harigaya Y, Parker R. 2016. Codon optimality and mRNA decay. Cell Res. 26: 1269–1270.

Hata H, Mitsui H, Liu H, Bai Y, Denis CL, Shimizu Y, Sakai A. 1998. Dhh1p, a putative RNA helicase, associates with the general transcription factors Pop2p and Ccr4p from Saccharomyces cerevisiae. Genetics 148: 571–579.

Ingolia NT. 2010. Genome-wide translational profiling by Ribosome footprinting. Methods Enzymol 470: 119–142.

Lim J, Lee M, Son A, Chang H, Kim VN. 2016. MTAIL-seq reveals dynamic poly(A) tail regulation in oocyte-to-embryo development. Genes Dev 30: 1671–1682.

Meyer S, Temme C, Wahle E. 2004. Messenger RNA turnover in eukaryotes: pathways and enzymes. Crit Rev Biochem Mol Biol 39: 197–216.

Miller C, Schwalb B, Maier K, Schulz D, Dümcke S, Zacher B, Mayer A, Sydow J, Marcinowski L, Dölken L, et al. 2011. Dynamic transcriptome analysis measures rates of mRNA synthesis and decay in yeast. Mol Syst Biol 7: 458.

Minvielle-Sebastia L, Winsor B, Bonneaud N, Lacroute F. 1991. Mutations in the yeast RNA14 and RNA15 genes result in an abnormal mRNA decay rate; sequence analysis reveals an RNA-binding domain in the RNA15 protein. Mol Cell Biol 11: 3075–3087.

Mishima Y, Tomari Y. 2016. Codon Usage and 3’ UTR Length Determine Maternal mRNA Stability in Zebrafish. Mol Cell 61: 874–885.

Muhlrad D, Parker R. 1992. Mutations affecting stability and deadenylation of the yeast MFA2 transcript. Genes Dev 6: 2100–2111.

Parker R. 2012. RNA degradation in Saccharomyces cerevisae. Genetics 191: 671–702.

Presnyak V, Alhusaini N, Chen YH, Martin S, Morris N, Kline N, Olson S, Weinberg D, Baker KE, Graveley BR, et al. 2015. Codon optimality is a major determinant of mRNA stability. Cell 160: 1111–1124.

Sheets MD, Fox CA, Hunt T, Vande Woude, Wickens M. 1994. The 3’-untranslated regions of c-mos and cyclin mRNAs stimulate translation by regulating cytoplasmic polyadenylation. Genes Dev 8: 926–938.

Sippel a E, Stavrianopoulos JG, Schutz G, Feigelson P. 1974. Translational properties of rabbit globin mRNA after specific removal of poly(A) with ribonuclease H. Proc Natl Acad Sci U S A 71: 4635–9.

Subtelny AO, Eichhorn SW, Chen GR, Sive H, Bartel DP. 2014. Poly(A)-tail profiling reveals an embryonic switch in translational control. Nature 508: 66–71.

Thore S, Mauxion F, Séraphin B, Suck D. 2003. X-ray structure and activity of the yeast Pop2 protein: a nuclease subunit of the mRNA deadenylase complex. EMBO Rep 4: 1150–1155.

Tucker M, Valencia-Sanchez MA, Staples RR, Chen J, Denis CL, Parker R. 2001. The transcription factor associated Ccr4 and Caf1 proteins are components of the major cytoplasmic mRNA deadenylase in Saccharomyces cerevisiae. Cell 104: 377–386.

Wang H, Morita M, Yang X, Suzuki T, Yang W, Wang J, Ito K, Wang Q, Zhao C, Bartlam M, et al. 2010. Crystal structure of the human CNOT6L nuclease domain reveals strict poly(A) substrate specificity. EMBO J 29: 2566–76.

Wiederhold K, Passmore L a. 2010. Cytoplasmic deadenylation: regulation of mRNA fate. Biochem Soc Trans 38: 1531–1536.

Yamashita A, Chang T-C, Yamashita Y, Zhu W, Zhong Z, Chen C-Y a, Shyu A-B. 2005. Concerted action of poly(A) nucleases and decapping enzyme in mammalian mRNA turnover. Nat Struct Mol Biol 12: 1054–1063.

Zheng D, Tian B. 2014. Sizing up the poly(A) tail: Insights from deep sequencing. Trends Biochem Sci 39: 255–257.

